# Histone deacetylase inhibitor givinostat attenuates nonalcoholic steatohepatitis and liver fibrosis

**DOI:** 10.1101/2020.06.09.141655

**Authors:** He-ming Huang, Shi-jie Fan, Xiao-ru Zhou, Yan-jun Liu, Xiao Li, Li-ping Liao, Jing Huang, Cui-cui Shi, Liang Yu, Rong Fu, Jian-gao Fan, Yuan-yuan Zhang, Cheng Luo, Guang-ming Li

## Abstract

Non-alcoholic steatohepatitis (NASH) is a common chronic liver disease that causes worldwide morbidity and mortality, yet there is still a lack of pharmacological therapies. Liver inflammation is an important contributor for disease progression from non-alcoholic fatty liver disease (NAFLD) to non-alcoholic steatohepatitis (NASH). We identified HDAC inhibitor givinostat as a potent inhibitor of macrophages inflammatory activation, and aimed to evaluate the therapeutic potential of givinostat for treatment of NASH. Daily administration of givoinostat (10mg/kg) alleviated inflammation and attenuated hepatic fibrosis in methionine- and choline-deficient diet (MCD)-induced NASH mice. RNA-seq analysis of liver tissues form MCD-fed mice revealed givinostat potently blocked expression of inflammation-related genes and regulated a broad set of lipid metabolism-related genes. In hepatocyte givinostat reduced palmitic acid induced intracellular lipid accumulation. The benefit of givinostat was further confirmed in fructose, palmitate, cholesterol diet (FPC) induced NASH mice. Givinostat attenuated hepatic steatosis, inflammation as well as liver injury in FPC-induced NASH. In conclusion, givinostat appears to be efficacious in reversing diet-induced NASH, and may serve as a therapeutic agent for treatment of human NASH.

## Introduction

Non-alcoholic fatty liver disease (NAFLD), a complicated liver metabolic disease, has become the most common chronic liver disease(1) and is considered as one of the leading causes of end-stage liver disease. The spectrum of NAFLD ranges from simple liver steatosis to non-alcoholic steatohepatitis (NASH). In western countries approximately one third population is potentially affected by NAFLD, which is related to obesity and type 2 diabetes mellitus (T2DM)(2). During the past few years, a growing number of people are suffering from NAFLD and the disease is increasingly prevalent worldwide. Modelling studies demonstrate that the prevalence of NAFLD will continue to rise in the next decade(3, 4). Progressive form of NAFLD, NASH, which is accompanied by liver injury and inflammation can progress to liver fibrosis, cirrhosis, and even hepatocellular carcinoma(HCC)(5). NASH has increasingly become an enormous clinical and economic burden, and cirrhosis and HCC caused by NASH will soon become a primary indication for liver transplantation(6), yet there is still a lack of efficient pharmacological therapies for the treatment of NASH.

Hepatic inflammatory response is a crucial driving force of disease progression, fueling the transition from NAFLD to NASH (7–9). Inflammation is not only central to the disruption of metabolic homeostasis in the liver but promotes sustained hepatic fibrogenesis and is present in virtually all patients with hepatic fibrosis. It also reports that patients’ inflammation level correlates with hepatic fibrosis progression(10) (11). Macrophage, including liver resident macrophages (Kupffer cells) and recruited monocyte-derived macrophages, are key players of hepatic inflammation. In NASH, free fatty acids (FFA) stimulate the production of a series of proinflammatory mediators including IL-1β, IL6, CCL5 in hepatocyte and commit macrophage towards an inflammatory activated state. The activation of macrophage further leads to hepatic stellate cell (HSC) activation and extracellular matrix (ECM) secretion and deposition. Macrophages can produce and activate the archetypal pro-fibrotic cytokine TGF-β and other soluble mediators which can act on the HSCs to induce a pro-fibrotic phenotype(12). Macrophage-HSC crosstalk is also mediated by IL-1 and TNFα, neutralization of which lead to decreased fibrosis in mouse models (13, 14). Previous animal studies and early clinical trials showed promising results specifically targeting macrophage. Depletion of macrophage by clodronate liposome or gadolinium chloride in high fat diet (HFD)-fed mice protected from the development of steatosis, and demonstrated decreased expression of inflammatory cytokines and fibrosis-related gene(15, 16). Pharmacological polarization of macrophage towards anti-inflammatory M2 phenotype partially reverse steatosis. Inhibition of CCL2/CCR2 or CCL5/CCR5, which target monocyte recruitment, has been shown to attenuate liver fibrosis(7, 17–19), and the CCR2/5 antagonist cenicriviroc is now in a phase 2 clinical trial(20). Therefore, targeting inflammation may help alleviate hepatic steatohepatitis and fibrosis and prevent the progression of NASH, thus representing a therapeutic strategy for treatment of NASH (21).

Epigenetics, an inheritable phenomenon that affects gene expression without altering the DNA sequence, provided a new perspective on the pathogenesis of NAFLD/NASH (22). Epigenetic alterations occurred during the adaption of hepatocytes to the lipotoxic environment, inflammation and oxidative stress, orchestrating the reprogramming of transcriptional machinery and contributing to the pathology of NASH (23). DNA methylation has not only been suggested to be involved in the progression of NAFLD towards advanced fibrotic clinical stages but also could affect NAFLD-related metabolic phenotypes, such as insulin resistance. Moreover, some epigenetic enzymes are starting to spring up in the biology background of NAFLD. For example, the histone methyltransferase Suv39h2 has been reported to contribute to NASH pathogenesis by promoting the inflammatory response in hepatocytes and macrophages. Suv39h2 deficiency in knockout mice protects the mice from NASH, thus Suv39h2 is considered as a potential target for the development of novel therapeutic solutions(24). Epigenetic machinery, has been shown to be highly involved in the regulation of inflammatory response(25, 26), yet its role in pathogenesis of NASH and whether drugs targeting epigenetic enzymes could be used for NASH treatment remains poorly studied. Given the crucial role of epigenetic machinery in regulation inflammation, which is associated with NASH progression, epigenetic inhibitors might show great potential in exploring and discovering novel therapies for treatment of NASH

Our study aimed to identify novel candidate compound that could alleviated inflammation and prevent NASH progression. We established a cell-based high-throughput assay based on inflammatory macrophages activation and screened our in-house epigenetic compound library (27). The HDAC inhibitor (HDACi) givinostat was identified as the most potent hit, which inhibited macrophages activation *in vitro*. In a methionine- and choline-deficient diet (MCD) mouse model, givinostat significantly alleviated hepatic inflammation and liver fibrosis. RNA-seq analysis of the liver tissue from MCD model revealed the gene affected by givinostat were involved in inflammation and lipid metabolism pathway. Further study on hepatocyte and fructose, palmitate, cholesterol diet (FPC) mouse model verified that givinostat could not only remit inflammation but also diminish intracellular lipid accumulation, making givinostat a promising drug candidate for developing new therapy for treatment of NASH.

## Methods

### Animals experiments

Male C57BL/6J 8- to 9-week-old mice (specific pathogen-free), with body weights ranging from 21 to 23g, were purchased from SIMM Animal Center (Shanghai, China). All animals were housed under standard laboratory conditions (21±2°C, 12 h light-dark cycle). All animal experiments were performed based on the institutional ethical guidelines on animal care and were approved by the Institute Animal Care and Use Committee at the Shanghai Institute of Materia Medica.

The animals were fed either a control diet or MCD diet (Jiangsu meidi bio-pharmaceutical co. LTD) ad libitum for 8 weeks. At the same time, MCD-diet mice were randomized to either the pharmacological HDACi givinostat (10mg/kg, formulated in PBS) daily or its vehicle solution by intraperitoneal (i.p.) injection for 8 weeks (n=10 per group). At the end of the experiment, the mice were anesthetized with 10% chloral hydrate, and their livers were immediately harvested.

The animals were fed either an FPC diet (Jiangsu meidi bio-pharmaceutical co. LTD) plus fructose 18.9g/l and glucose 23.1g/l add to the drinking water, or control diet ad libitum for 16 weeks. After 6 weeks on the diet, FPC-fed mice were randomized to either givinostat (10mg/kg, formulated in PBS) daily or its vehicle solution by i.p. injection for 10 weeks (n=10 per group). At the end of the experiment, the mice were anesthetized with 10% chloral hydrate, and their livers were immediately harvested.

### Histology and immunohistochemical analysis

Liver histology was performed using fixed in 4% paraformaldehyde, embedded in paraffin, sectioned, and stained with hematoxylin and eosin (H&E; Wuhan goodbio G1005) and Sirius red stain (Wuhan goodbio G1018). For immunohistochemistry (IHC), paraformaldehyde-fixed paraffin-embedded liver tissue sections were deparaffinized, hydrated, and stained with antibody against F4/80 (1:100, MAB5580,RD), CD68 (1:100, MAB0803, Novus), α-SMA (1:100, GB13044, RD) and Col 1a1 (1:100, GB11022-1, RD). Subsequently, the slides were further processed using corresponding secondary antibodies, followed by counterstained with hematoxylin. Sirius red stained areas and F4/80, CD68, α-SMA and Col 1a1 immunopositive areas were quantified by digital image analysis of 10 random fields per slide using the Image-Pro Plus (Media Cybernetic, Inc.). Lipid droplet accumulation in the liver was observed using Oil red O (O0625; sigma-Aldrich) staining of frozen sections, frozen liver sections stained with Oil red O staining reagent for 30 min and were counterstained with hematoxylin. To calculate the NAS score, H&E images were analyzed according to criteria described by Brunt et al(28). The NAS score consists of three components: steatosis, lobular inflammation and hepatocyte ballooning.

### Cell culture

The mouse macrophage cell line RAW264.7 was cultured in Dulbecco’s Modified Eagle Medium (DMEM) containing 10% heat-inactivated fetal bovine serum (FBS, Gibco, Australia) at 37°C, 5% CO2 with complete media. For stimulation, the cells were treated with LPS (Escherichia coli, 055:B5, Sigma, 1μg/ml) and/or givinostat (9μM, 3μM, 1μM) for 4h or 24h. For lipid stimulation, the cells were treated with palmitate (PA) (0.4mM, P9767, Sigma-Aldrich) and/or givinostat (4μM, 1μM) for 12 h.

The human hepatocellular carcinoma cell line HepG2 was cultured in DMEM containing 10% FBS at 37°C, 5% CO2 with complete media. To mimic the *in vitro* hepatic steatosis model, the cells were treated with PA (0.4mM) and or givinostat (9μM, 3μM, 1μM) for 12 h.

### RNA-Seq analysis

Total RNA isolated from liver tissues form control mice, vehicle treated MCD mice and givinostat-treated MCD mice. RNA Integrity Number (RIN) value was used to assess the quality of the isolated RNAs. Only RNAs with RIN≥7.0 were used for sequencing. The sequencing reads were located to mm10 by STAR 2.5 and the gene counting was quantified using feature counting software. The Deseq2 R package was used for differential gene expression analysis. The p value was adjusted by the Benjamini and Hochberg methods, and the 5% FDR cutoff value and the fold change greater than 1.5 were set as the threshold of the significant gene. The differentially expressed genes were further analyzed by gene-annotation enrichment analysis using the KOBAS3.0 bioinformatics platform.

### RNA extraction and RT-qPCR

Total RNA was isolated with the total RNA extraction reagent (Vazyme, China) form liver tissues and cells, and then reverse transcribed with the special cDNA synthesis kit (Vazyme, China) according to the manufacturer’s protocol. Quantification of gene expression was measured by RT-qPCR on Quant Studio 6 Flex Real -Time PCR system (ABI). Target genes expression was calculated using the ∆∆Ct method and expression was normalized with GAPDH expression levels. The primer sequences are listed in Supplemental Table1.

### Western blotting

Total protein was extracted from RAW264.7 cells and frozen liver tissues with RIPA lysis buffer. BCA Protein Assay Kit (Sangon Biotech, China) was used to measured protein concentrations. Protein samples were separated by sodium dodecyl sulfate (SDS)-PAGE and transferred onto a nitrocellulose membrane (Millipore, Temecula, CA, USA). After blocking with 5% non-fat powdered milk, the nitrocellulose membrane was incubated with primary antibodies at 4°C overnight, followed by incubation with the corresponding secondary antibodies for 1h at room temperature. A ChemiScope3400 imaging system was used for signal detection. Protein expression levels were quantified by ImageJ and normalized to levels of GAPDH. The following primary antibodies were used: anti-α-SMA (Cell Signaling Technology, 19245S, 1:1000), anti-Col1a1 (Cell Signaling Technology, 39952S, 1:1000), anti-H3 (Cell Signaling Technology, 4499S, 1:1000), anti-ac-lysine (Cell Signaling Technology, 6952S, 1:1000), and GAPDH (Cell Signaling Technology, 5174S, 1:5000).

### Biochemical analysis

Serum aspartate aminotransferase (AST), alanine aminotransferase (ALT) and alkaline phosphatase (ALP) were assessed using a Hitachi 7020 automatic analyzer (Hitachi, Tokyo, Japan).

### ELISA

The concentration of IL-1β, IL-6 and TNF-α in RAW264.7 cells culture supernatants were analyzed using ELISA kits (No. EMIL6RA, 88-7324-22, bms6002; Thermo Scientific) according to the manufacturer’s instructions.

### Liver TG, TC and intracellular TG levels quantification

Liver TG and TC levels were measured from mouse liver homogenates. Briefly, for TG quantification, wet liver tissue(100mg) or cells (1*10^7^) were homogenized in a 5% NP-40 solution. Triglyceride Quantification Kit (Sigma-Aldrich, MAK266) was used for the assay according to manufacturer’s instructions. Photometric absorbance was read at 570nm using microplate reader (Thermo scientific, Multiskan FC). For liver TC quantification, 10mg wet liver tissue were homogenized in a chloroform: isopropanol: IGEPAL CA-630 solution. Cholesterol Quantitation Kit (Sigma-Aldrich, MAK043) was used for the assay according to manufacture’s instruction. Photometric absorbance was read at 570nm using microplate reader (Thermo scientific, Multiskan FC).

### Statistical analysis

All results were expressed as mean ± S.D., and the differences between groups were performed using GraphPad Prism 7.0 statistical software (GraphPad Software, Inc., La Jolla, CA, USA). P<0.05 was considered a significance.

## Result

### Givinostat inhibited LPS and PA-induced inflammation *in vitro* and alleviated MCD-induced liver inflammation *in vivo*

Our previous high-throughput screening identified givinostat as one of the most potent inhibitors that reduced macrophage inflammatory activation. Given the crucial role of inflammation in NASH progression, we aimed to test whether givinostat could reduce liver inflammation and fibrosis, and evaluate its therapeutic efficacy for treatment of NASH. We first confirmed its anti-inflammatory effects *in vitro*. RAW264.7 cells were stimulated by lipopolysaccharide (LPS) in the presence of givinostat to examine effect of givinostat on LPS-induced cytokine expression, which is indicative of macrophage inflammatory activation. We cultured RAW264.7 cells in the presence or absence of givinostat and treated them with LPS(1ug/ml) for 4 or 24 h. RT-qPCR was used to detect the mRNA expression levels of IL-6, IL-1β and TNF-α, which are representative inflammatory mediators that correlated with severity of inflammation response. Data showed that LPS induced a significant increase in the mRNA and protein expression levels of IL-6, IL-1β and TNF-α (Figure 1A 1B). Co-treatment with givinostat abrogated the LPS-induced transcription of these cytokines in a concentration-dependent manner (Figure 1A). Consistently, givinostat treatment decreased the protein expression and secretion of IL-6, IL-1β and TNF-α induced by LPS stimulation (Figure 1B), as shown by ELISA analysis of cell supernatant. When exposed to palmitic acid (PA), RAW264.7 cells up-regulated the production of proinflammatory mediators including IL6, IL-1β and TNF-α. Givinostat significantly dampened PA-induced transcription of proinflammatory mediators (Figure 1C). Notably, givinostat administration increased protein level of acetylated Histone 3 and 4 in RAW264.7 (Figure 1D), validating its on-target inhibition of HDAC.

**Figure1.**
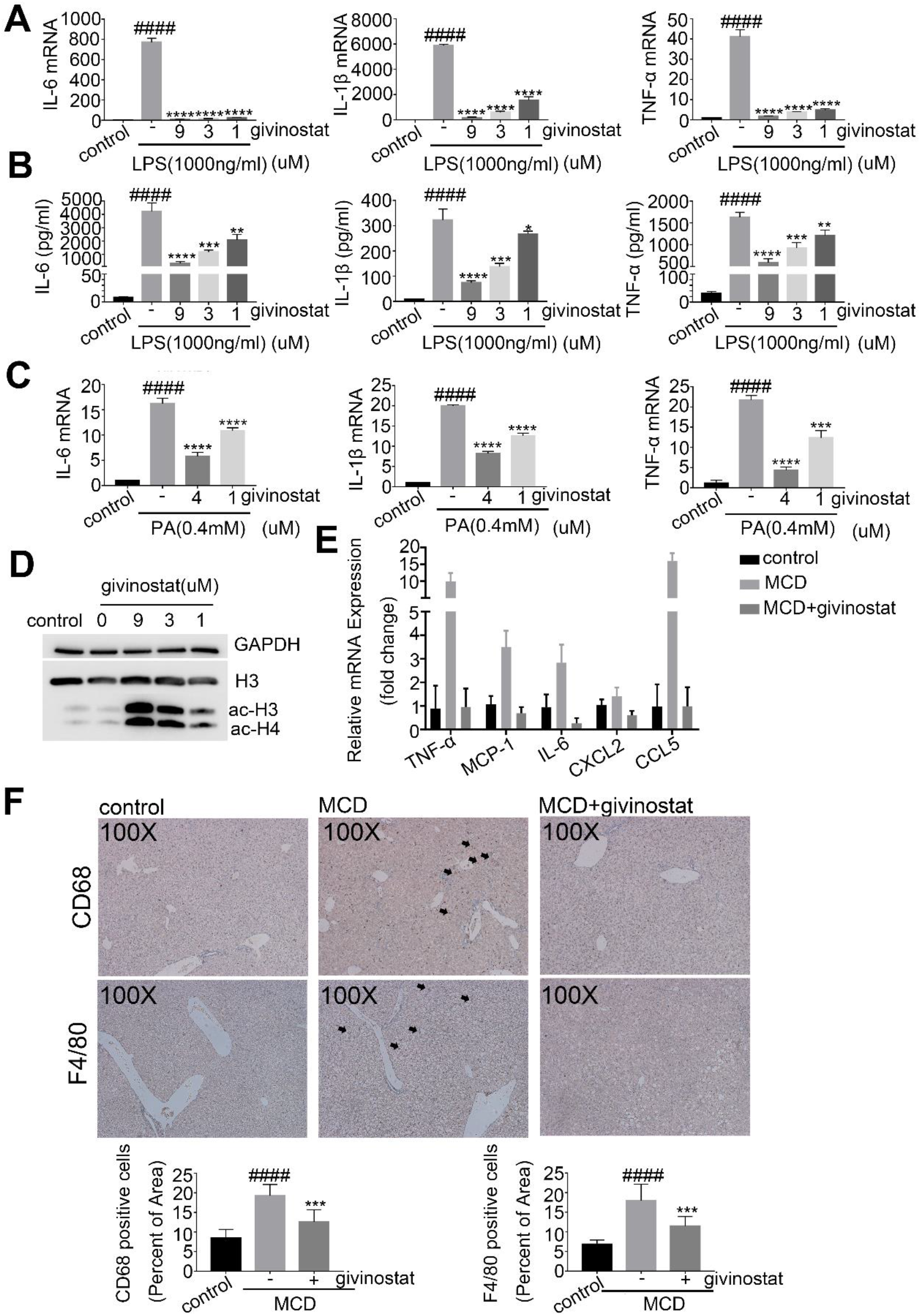
Givinostat inhibited LPS and PA-induced inflammation *in vitro* and alleviated MCD-induced liver inflammation *in vivo*. (A)RAW264.7 cells were treated with LPS and givinostat for 4h, after which the IL-6, IL-1β and TNF-α mRNAs were quantified by RT-qPCR analysis. (B) RAW264.7 cells cells were treated with PA an d givinostat for 12h, after which the IL-6, IL-1β and TNF-α mRNA were quantified by RT-qPCR analysis. (D RAW264.7 cells were treated with LPS and givinostat for 4h the acetylated H3 and H4 protein levels were quantified by western blot analysis. (E) Relative mRNA levels for genes involved in inflammation in the liver of mice were fed either chow or MCD diet for 8 weeks, and givinostat or vehicle was given simultaneously. (n=10 mice per group). (F) Representative F4/80 and CD68 IHC staining and their quantification in the liver of mice were fed either chow or MCD diet for 8 weeks, and givinostat or vehicle was given simultaneously. (n=10 samples from 10 mice per Bars represent mean ± SD. ####P < 0.0001 versus control or control diet fed mice. **P < 0.01, ***P < 0.001,0.001, ****P < 0.0001 versus LPS or PA group or vehicle treated MCD fed mice.

Given the dramatic reduction of inflammation by givinostat we observed *in vitro*, we examined whether givinostat could also alleviate hepatic inflammation *in vivo* in MCD-fed mice. To determine whether givinostat affected inflammation, we examined the expression of cytokines and chemokines in livers from both givinostat-treated and vehicle-treated mice. In the livers of MCD-fed mice, concentrations of the inflammatory mediators including IL-6, TNF-α, MCP-1 (also known as CCL2), CCL5 and CXCL2 were significantly higher compare to control diet-fed mice(Figure 1E). MCD-fed mice were treated with either givinostat (10mg/kg) daily by intraperitoneal (i.p.) injection or its vehicle. Expression of these inflammatory cytokines in the livers of givinostat-treated mice was reduced in comparison to that of vehicle-treated mice after 8 weeks on MCD diet (Figure 1E). During hepatic inflammation, activated macrophage secreted CCL2 and CCL5, which recruit monocyte to the liver and further exaggerate/deteriorate liver inflammation. Recruitment of monocyte and its differentiation into inflammatory macrophage is an important step for NASH progression. The inhibition of CCL2 and CCL5 expression by givinostat *in vitro* and *in vivo* prompted us to examine whether givinostat intervention could inhibit macrophages infiltration in liver of MCD mice by performing immunohistochemical (IHC) staining of liver tissues. Results showed that macrophages marked by F4/80 and CD68 increased in the liver of MCD diet-fed mice compared with the control mice, and was significantly reduced after givinostat treatment (Figure 1F). Taken together, these data demonstrated that givinostat treatment significantly reduced liver inflammation and infiltration of inflammatory cells *in vivo* in MCD-fed mice.

### Givinostat can protect against MCD-induced liver fibrosis

The liver inflammation is an important driving force of disease progression, as it promotes sustained liver fibrogenesis. We next examined whether givinostat could also alleviated liver fibrosis in MCD-fed mice. Sirius Red staining is commonly used to visually assess collagen levels associated with liver fibrosis. MCD-fed mice developed significant liver fibrosis as demonstrated by increased collagen deposition measured by Sirius Red staining, whilst, givinostat-treated mice showed impressively reduced amounts of collagen fibers compared with vehicle-treated mice(Figure 2A). Morphometric analysis yielded concordant results where the Sirius Red stained collagen areas were significantly reduced in givinostat-treated mice compared to vehicle-treated mice (P<0.05) (Figure 2A). Consistently, IHC staining of α-SMA and collagen type1(Col 1a1), the most abundant ECM protein in the fibrotic liver tissue, demonstrated that givinostat notably attenuated increased expression of α-SMA and Col 1a1 in the liver of MCD-fed mice compared to vehicle-treated mice (Figure 2B). Consistently, givinostat reduced mRNA expression of Col1a1 and α-SMA in the liver of MCD-fed mice (Figure 2C). Furthermore, the liver Col1a1 and α-SMA protein expression were also markedly reduced compared with that in vehicle-treated MCD-fed mice (Figure 2D). Collectively, these data indicate that givinostat significantly alleviated liver fibrosis in the MCD-fed mice.

**Figure2.**
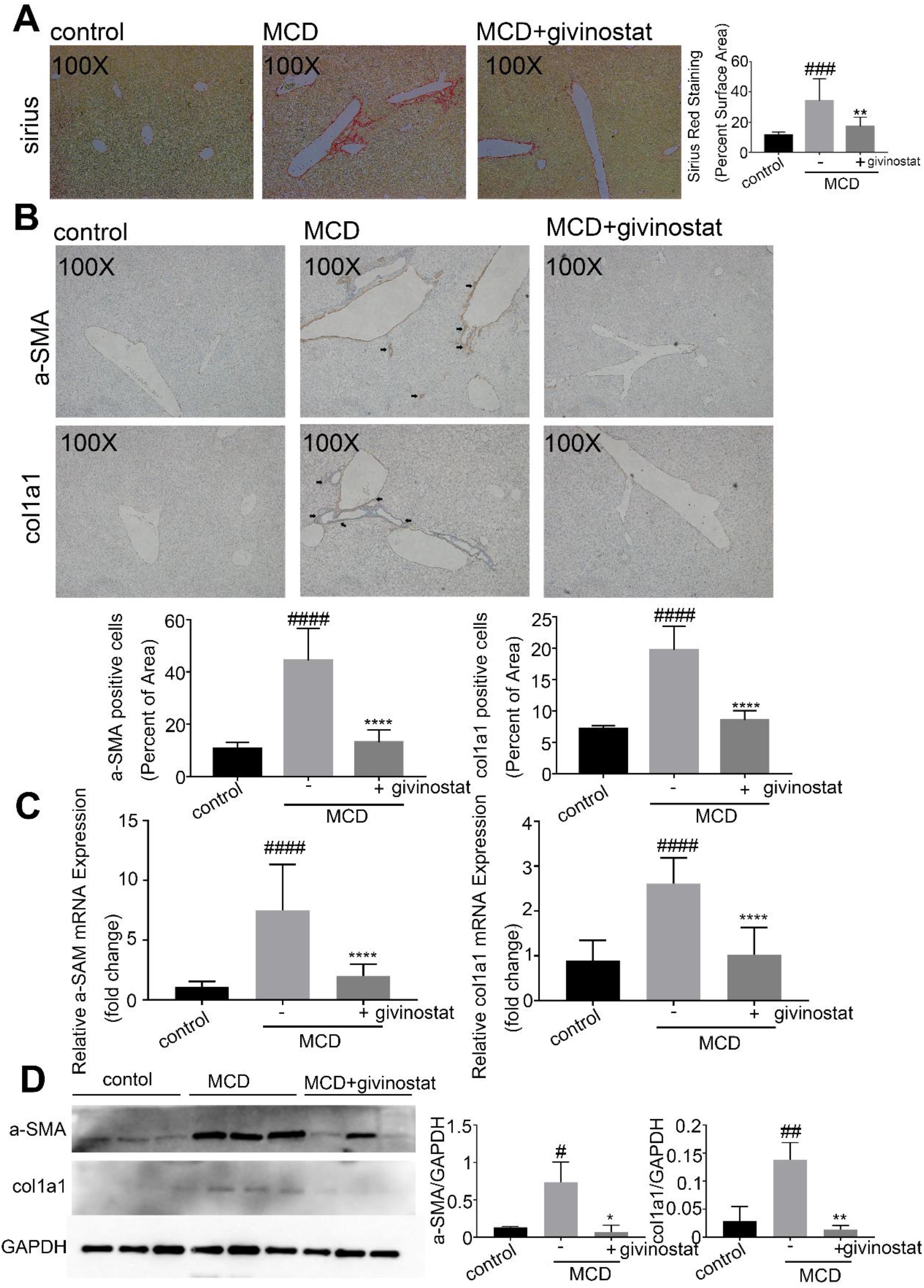
Givinostat can protect against MCD-induced liver fibrosis. C57BL/6J mice were fed either chow or MCD diet for 8 weeks, and givinostat or vehicle was given simultaneously. (A). Collagen deposition by Sirius Red staining of livers sections and their quantification shown in the right panel (n=10 samples from 10 mice per group) (B). Representative Col1a1 and α-SMA IHC staining, and their quantification shown in the lower panel (n=10 samples from 10 mice per group). (C). Representative RT-qPCR analysis of liver Col1a1 and α-SMA mRNA expression. (D). Representative western blot analysis of the liver Col1a1 and α-SMA protein expression, with quantification shown in the right panel. Bars represent mean ± SD. #P<0.05, ##P<0.01, ###P<0.001, ####P<0.0001 versus control diet-fed mice. *P<0.05, **P<0.01, ***P<0.001, ****P<0.0001 versus vehicle-treated MCD-fed mice.

### Givinostat regulated inflammatory and metabolism gene expression *in vivo* as determined by RNA-Seq analysis

Next we investigated into the mechanism underlying the therapeutic efficacy of givinostat. RNA sequencing (RNA-seq) analysis was performed to compare the gene expression profiles of liver tissues from control mice, vehicle treated MCD mice and givinostat-treated MCD mice, followed by principle component analysis (PCA) and differential expression genes (DEGs) analysis. Then DEGs were analyzed by KEGG pathway enrichment. In PCA, principle component 1 (PC1) and principle component 2 (PC2) take a high proportion of variance, which means they can explain the difference between groups (Figure 3A). Based on PC1 and PC2, three groups are classified distinctly, which means that givinostat treatment could change MCD condition (Figure 3B). Then, we analyzed DEGs between Model group and HDACi givinostat-treated group. Then DEGs defined as p.adj < 0.05 and log2FC >= 1.5 were selected to undergo subsequent KEGG pathway enrichment.

**Figue3.**
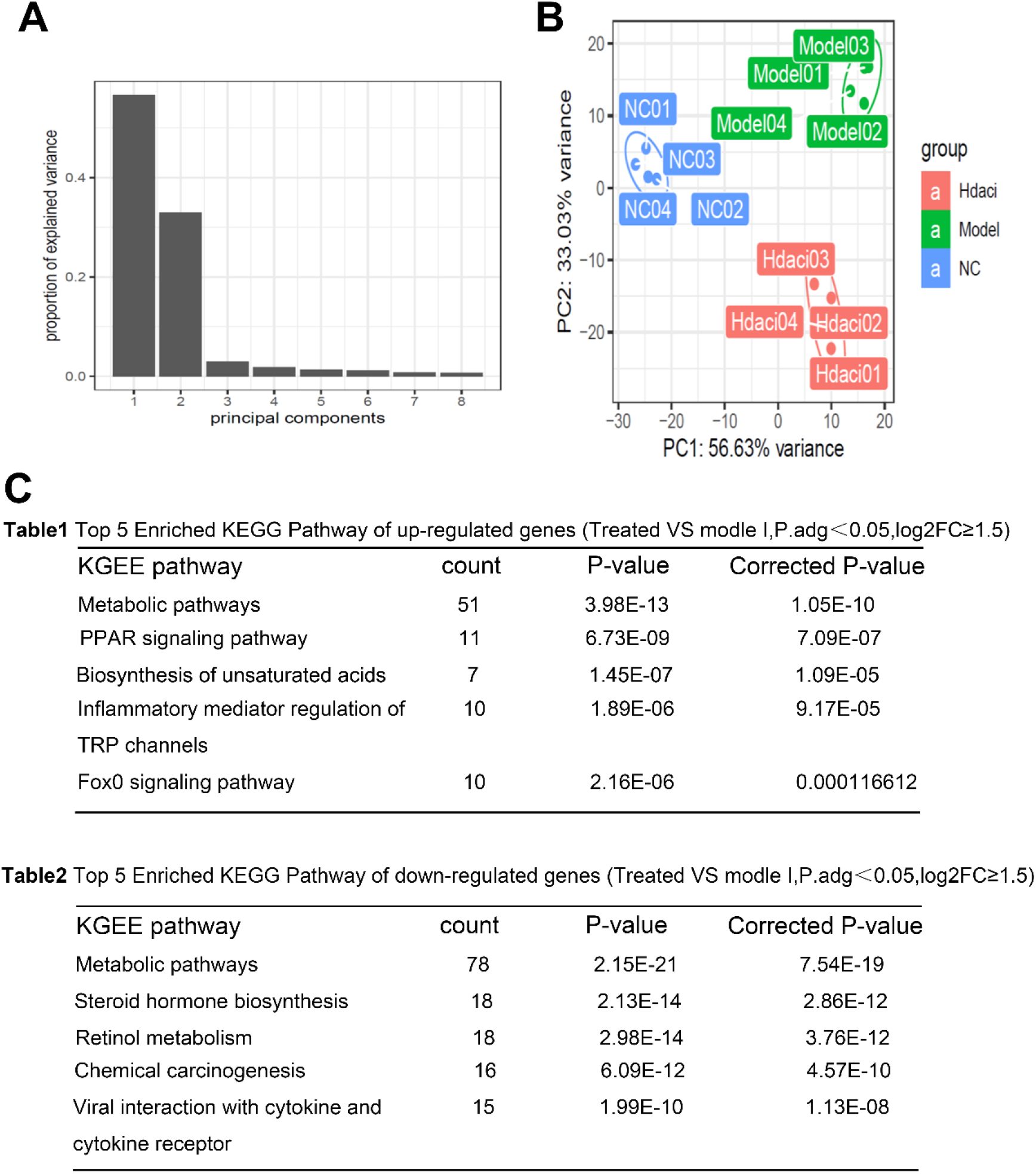
Givinostat regulated inflammatory and metabolism gene expression *in vivo* as determined by RNA-Seq analysis. RNA-seq analysis was performed on liver tissues extracted from mice fed either chow or MCD diet for 8 weeks, and givinostat or vehicle was given simultaneously. (n=4 mice per group). (A-B). Principle component analysis among three groups. (C). KEGG pathway analysis using differentially expressed genes between vehicle-treated and givinostat-treated MCD-fed mice, showing that the most significantly enriched pathway.

Consistent with above RT-qPCR data, givinostat shows its inflammation modulating effect because several inflammatory pathways are regulated. TRP channels pathway and FoxO signaling pathway which is important in inflammatory mediator regulation and immune cell regulation are enriched in up-regulated genes. Viral interaction with cytokine and cytokine receptor pathway is enriched in down-regulated genes, in agreement with CXCL2, CCL5 down-regulation observed in RT-qPCR.

Intriguingly, we found many genes related to metabolism regulation enriched. In both up or down regulated genes, “Metabolic pathways” ranks first and some other important metabolism pathways like PPAR, steroid biosynthesis, and unsaturated fatty acid biosynthesis follow closely (Figure 3C). From RNA-seq results, we conclude that givinostat potently blocked pathologic pro-inflammatory genes and regulated a broad set of lipid metabolism-related genes, prompting us to explore the protective effect of givinostat on hepatic steatosis

### Givinostat alleviate MCD-induced hepatic steatosis

We further elucidated the effects of givinostat on hepatic steatosis in MCD fed mice. The liver tissue of MCD diet exhibited obvious steatosis, ballooning hepatocytes, scattered lobular inflammatory cell infiltration and inflammatory foci, as indicated by HE and Oil red O staining (Figure 4A, 4B, middle panel). In contrast, the givinostat-treated mice showed minimal steatosis, with only a few small foci of steatosis observed in the centrilobular area (Figure 4A,4B, right panel). NASH scores further confirmed that givinostat treatment resulted in a decrease in hepatic steatosis, ballooning of hepatocytes and inflammation compared to vehicle-treated mice (Figure 4C). Consistently, the livers of vehicle-treated MCD-fed mice exhibited significantly increased lipid accumulation and content compared with control-fed mice, whereas the livers of givinostat-treated MCD-fed mice showed significantly reduced liver triglyceride (TG) and cholesterol (TC) levels (Figure 4D). Moreover, RT-qPCR analysis of liver samples indicated that the mRNA expression of genes involved in fatty acid β-oxidation (cholesterol 7α-hydroxylase [*CYP7A*], acyl-coenzyme A oxidase 1 [*ACOX-1*], Cytochrome c oxidase subunit 5A [*COX5A*], uncoupling protein 2 [*UCP2*] and pyruvate dehydrogenase kinase4 [*PDK4*]) were greatly attenuated, whereas genes involved in cholesterol & fatty acid synthesis (Fatty acid synthase [*FAS*], sterolregulatory element-binding protein-1c [*SREBP-1c*] and stearoyl-CoA desaturase-1 [*SCD1*]) were significantly increased in vehicle-treated MCD-fed mice compared to control fed mice (Figure 4E). Givinostat-treated MCD-fed mice exhibited an opposite pattern of expression of these lipid metabolic genes in the liver compared to vehicle-treated MCD-fed mice(Figure 4E). In addition, the plasma levels of alanine aminotransferase (ALT) and aspartate aminotransferase (AST), major markers of liver function, were elevated in vehicle-treated MCD-fed mice compared with control fed mice. Givinostat treatment significantly reduced serum AST and ALT activity in MCD-fed mice (Figure 4F), suggesting alleviated liver damage. Together, these findings demonstrate that givinostat treatment inhibited hepatic hepatosis, inflammation as well as fibrosis in MCD mice, thus improving NASH score and alleviating liver injury.

**Figure4.**
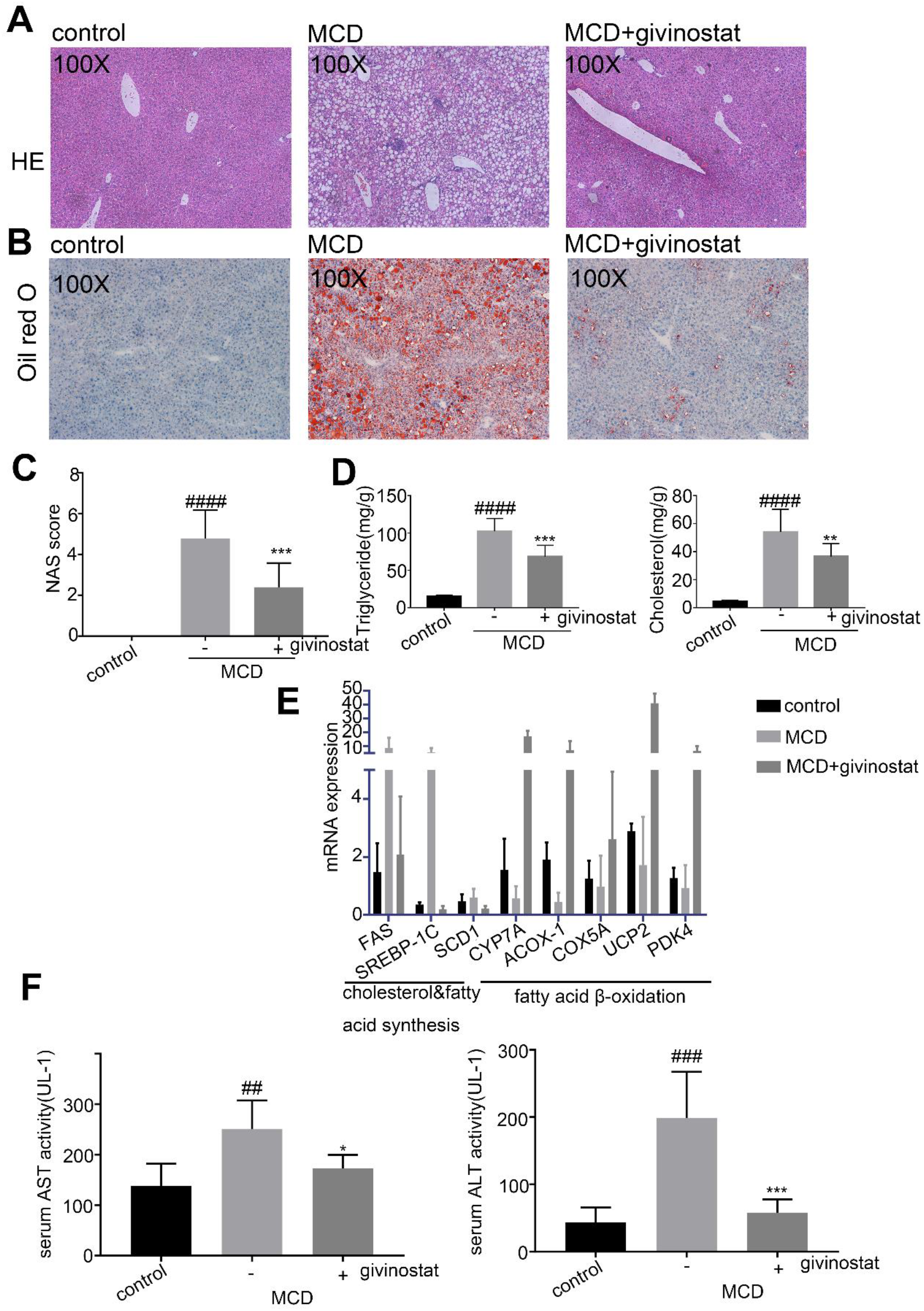
Givinostat alleviate MCD-induced hepatic steatosis. C57BL/6J mice were fed either chow or MCD diet for 8 weeks, and givinostat or vehicle was given simultaneously. (A). Representative images of HE stained liver sections. (B). Representative images of Oil Red O stained liver sections. (C). The NASH scores (n=10 mice per group). (D). The TG and TC levels in the livers (n=10 mice per group). (E). Transcript levels of lipid metabolism-related genes in the livers (n=10 mice per group). (F). The plasma ALT and AST levels in mice. (n=10 mice per group). Bars represent mean ± SD. ##P<0.01, ###P<0.001, ####P<0.0001 versus control diet-fed mice. *P < 0.05, ***P < 0.001, ****P < 0.0001 versus vehicle-treated MCD-fed mice.

### Givinostat diminish PA induced intracellular lipid accumulation in hepatocytes

Givinostat alleviated hepatic steatosis and regulated a broad set of lipid metabolic genes. We next examined whether givinostat directly affected lipid accumulation in hepatocyte *in vitro*. We used a PA-induced *in vitro* fatty liver cell model. We induced lipid accumulation in hepatic cells by exposing them to PA, to simulate excessive influx of free fatty acids into hepatocytes. The experiments were performed using PA complexed with bovine serum albumin (BSA) at a FFA to BSA ratio of 5.2:1(29).

Human hepatoma cells (HepG2) were cultured with 0.4 mM PA for 12h in the presence or absence of givinostat, as a control, HepG2 cells were treated with equimolar BSA carrier protein. HepG2 cells treated with 0.4mM PA for 12h exhibited significant increased lipid droplet accumulation compared to control cells treated with BSA, while minimal staining for lipids was seen in givinostat-treated cells, as indicated by Oil red O staining (Figure 5A). The availability of FFA increases with increased intake and is expected to enhance liver TG synthesis. We measured intracellular TG content in HepG2 cells that were exposed to 0.4mM PA for 12h (Figure 5B). Compared with the BSA control cells, PA-treated cells exhibited significantly increased lipid accumulation and content, whereas the givinostat-treated cells presented the opposite trend compared with PA-treated cells, as indicated by TG levels (Figure 5B). Thus, we observed a direct inhibitory effect of givinostat on lipid accumulation *in vitro*.

**Figure5.**
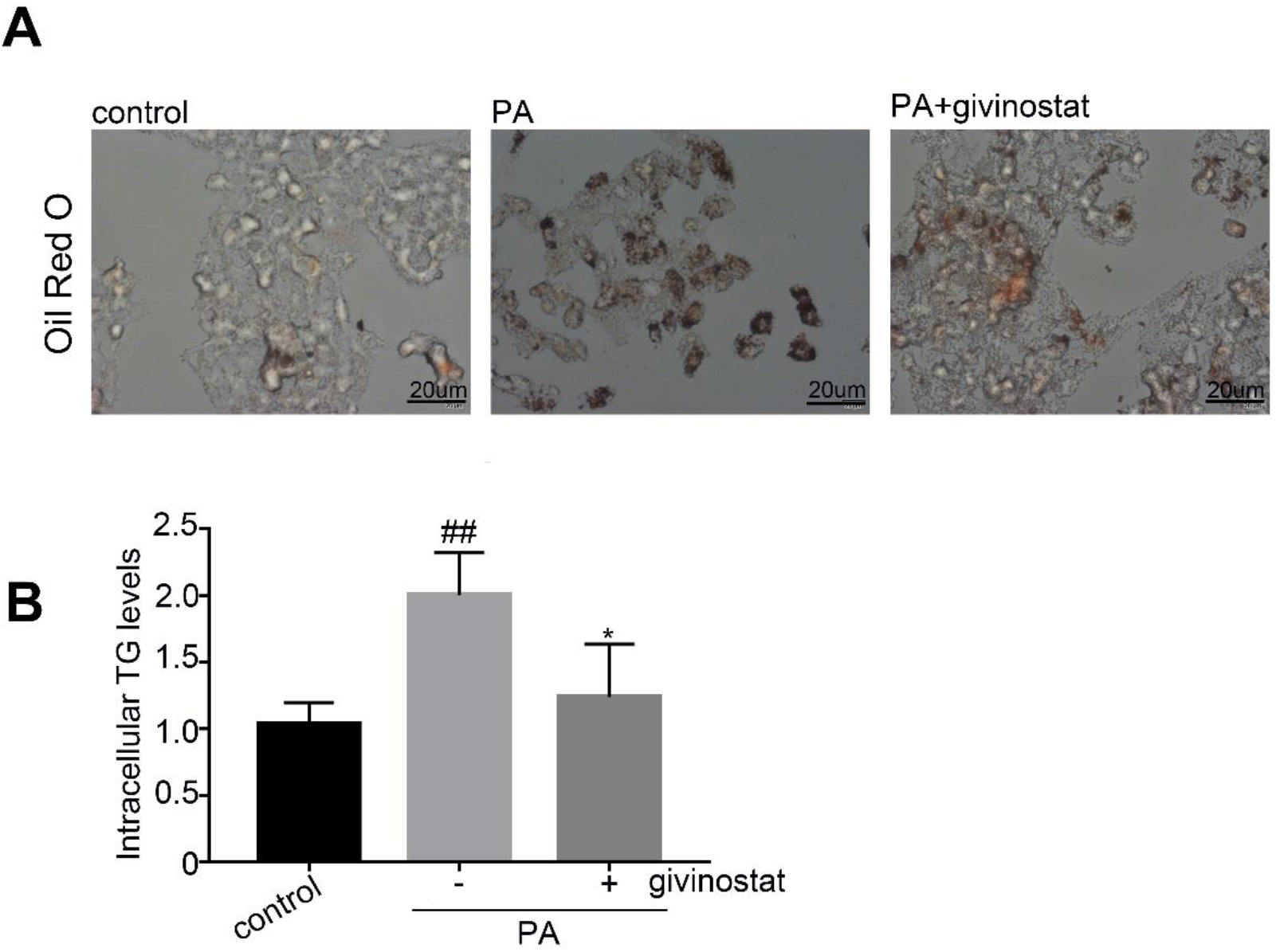
Givinostat diminish palmitic acid induced intracellular lipid accumulation in hepatocytes. (A). Representative images of Oil Red O staining of HepG2 cells that were treated with vehicle (control) or PA for 12h. (B). The intracellular TG content in HepG2 cells after treatment with PA or vehicle for 12h. Bars represent mean ± SD. ##P<0.01, ###P < 0.001, ####P < 0.0001 versus BSA control cells. *P < 0.05, ***P < 0.001, ****P<0.0001 versus PA treated cells.

### FPC-induced liver inflammation is diminished by givinostat

Encouraged by the potent regulatory effects of givinostat on hepatocyte metabolism and inflammation, we assessed the therapeutic efficiency of givinostat for NASH treatment and examined its impact on liver inflammation and lipid metabolism using mouse NASH models induced by diet rich in fructose, palmitate, cholesterol (FPC)(30). C57BL/6J mice were fed either chow or FPC diet for 16 weeks, and givinostat or vehicle was given during the last 10 weeks. After 16 weeks, FPC-fed mice had significantly higher serum ALT, AST and alkaline phosphatase (ALP) levels when compared with control diet fed mice, whereas givinostat-treated FPC-fed mice had a considerable reduction of serum ALT, AST and ALP levels compared with vehicle-treated FPC-fed mice, reflecting reduced liver injury (Figure 6A). IHC staining of liver tissues showed that vehicle-treated FPC-fed mice had a significant increase in macrophage (marked by F4/80 and CD68) in the liver than in liver of control diet fed mice, which was remarkably reduced by givinostat treatment (Figure 6B). A significant increase in mRNA levels of inflammation genes, including IL-6, TNF-α, MCP-1, CCL5 and CXCL2, was observed in livers of vehicle-treated FPC-fed mice compared to control diet fed mice (Figure 6C). Relative to FPC-fed mice treated with vehicle, givinostat-treated mice exhibited a considerable reduction in inflammatory gene expression in the liver (Figure 6C). Taken together, these data demonstrate givinostat treatment reduced liver dysfunction and inflammatory response in FPC-diet induced NASH mice model.

**Figure6.**
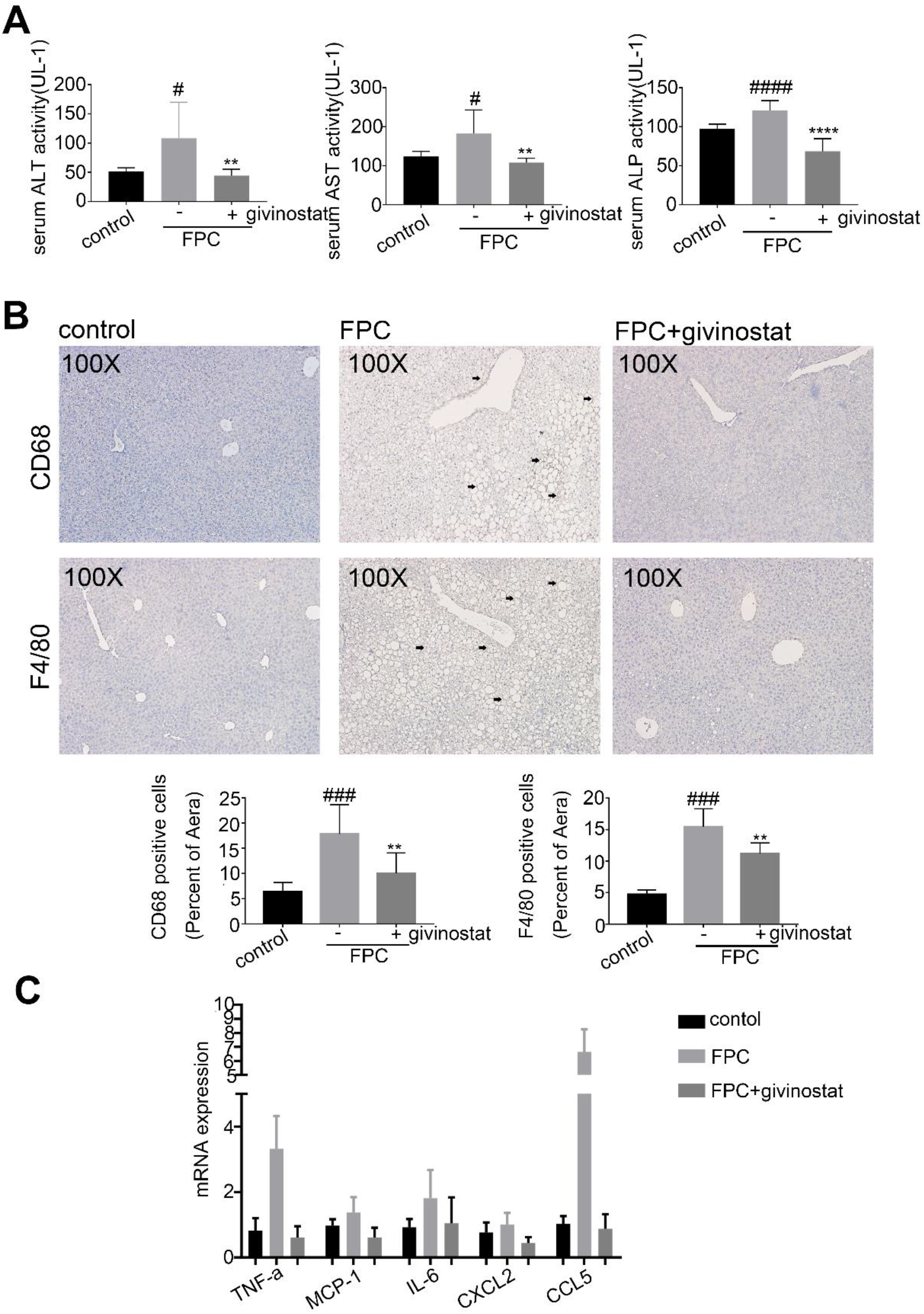
FPC-induced liver inflammation is diminished by givinostat. C57BL/6J mice were fed either chow or FPC diet for 16 weeks, and givinostat or vehicle was given during the last 10 weeks. (A). The plasma ALT, AST and ALP levels in mice (n=10 mice per group). (B). Representative F4/80 and CD68 IHC staining and their quantification (n=10 samples from 10 mice per group). (D). Relative mRNA levels for genes involved in inflammation in the liver (n=10 mice per group). Bars represent mean ± SD. #P<0.05, ##P<0.01, ####P<0.0001 versus control diet fed mice. **P<0.01, ***P<0.001, ****P<0.0001 versus vehicle-treated FPC-fed mice.

### FPC-induced hepatic steatosis is diminished by givinostat

We further examined whether givinostat could ameliorate steatosis in FPC-fed NASH mice. Vehicle-treated FPC-fed mice, in comparison to control diet fed mice, exhibited strikingly greater levels of steatosis, ballooning hepatocytes, scattered lobular inflammatory cell infiltration and inflammatory foci in the liver, as indicated by HE and Oil red O staining, which were reversed by givinostat treatment (Figure 7A,7B). Consistently, givinostat treatment significantly reduced the NASH scores in FPC-fed mice (Figure 7C).Furthermore, the livers of vehicle-treated FPC-fed mice exhibited significantly increased lipid accumulation and content compared with control-fed mice, whereas the livers of givinostat-treated FPC-fed mice presented the opposite trend compared with vehicle-treated FPC-fed mice, as indicated by liver TG and TC levels (Figure 7D). Moreover, RT-qPCR analysis of liver samples indicated that the mRNA expression of genes involved in fatty acid β-oxidation (*CYP7A*, *ACOX-1*, *COX5A*, *UCP2* and *PDK4*) were greatly attenuated, whereas cholesterol & fatty acid synthesis (*FAS*, *SREBP-1c* and *SCD1*) were significantly increased in vehicle-treated FPC-fed mice compared to control fed mice (Figure 7E). However, givinostat-treated FPC-fed mice exhibited a opposite pattern of expression of these lipid metabolic genes in the liver compared to vehicle-treated FPC-fed mice. Together, givinostat resulted in remarkable reduction in hepatic inflammation, lipid accumulation and liver dysfunction, as well as a significant improvement in fatty acid synthesis and oxidation in FPC-fed mice.

**Figure7.**
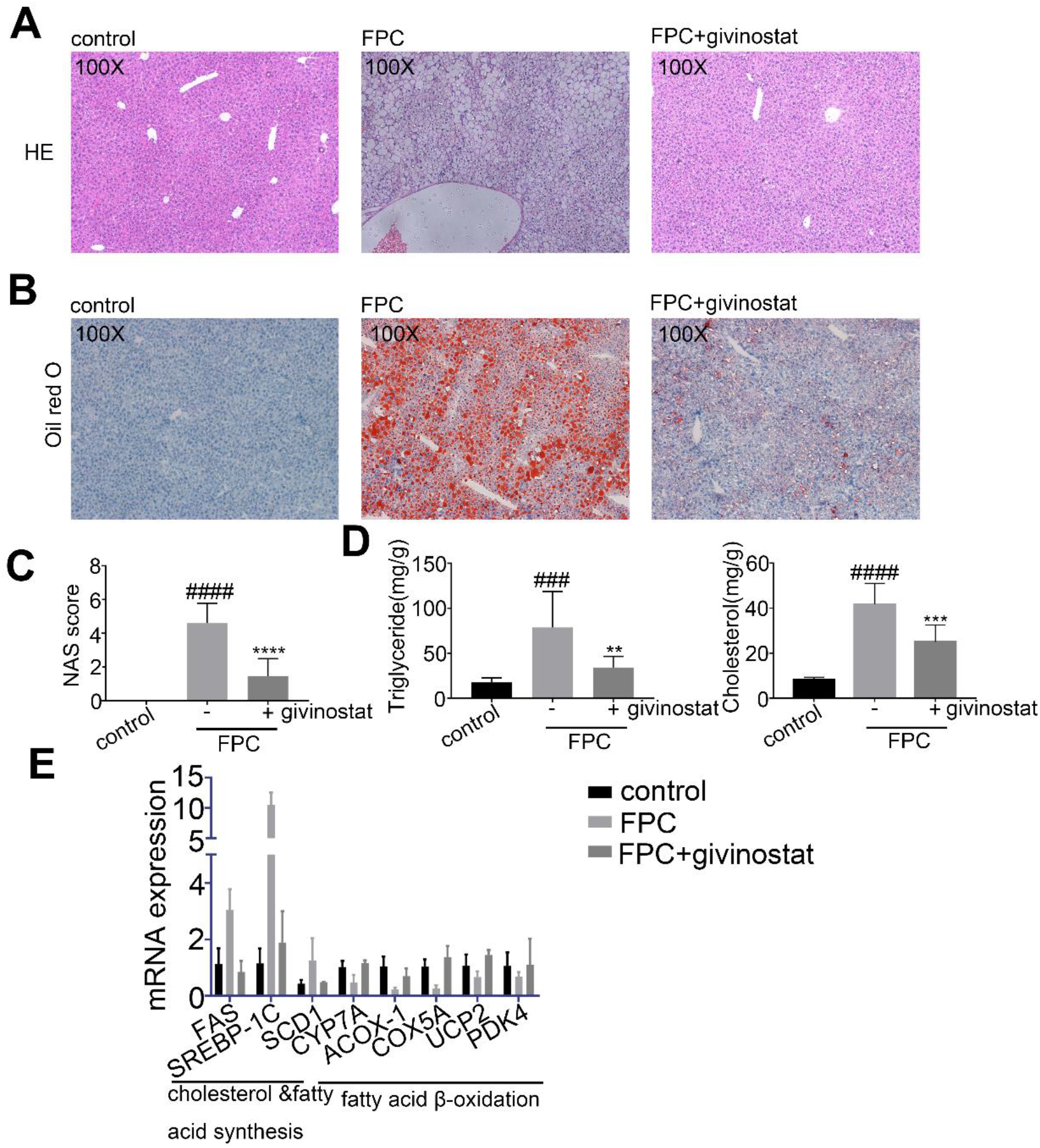
FPC-induced hepatic steatosis is diminished by givinostat. C57BL/6J mice were fed either chow or FPC diet for 16 weeks, and givinostat or vehicle was given during the last 10 weeks.(A). Representative images of HE stained liver sections (B). Representative images of Oil Red O stained liver sections. (C). The NASH scores (n=10 mice per group). (D). The TG and TC levels in the livers (n=10 mice per group). (E). Transcript levels of lipid metabolism-related genes in the livers (n=10 mice per group). Bars represent mean ± SD. ##P<0.01, ###P<0.001, ####P < 0.0001 versus control diet-fed mice. *P < 0.05, ***P < 0.001, ****P < 0.0001 versus vehicle-treated FPC-fed mice.

## Discussion

NAFLD is emerging as a worldwide public health threat(31). A number of patients with NAFLD develop a more inflammatory and progressive subtype-NASH, which can lead to cirrhosis and end-edge liver disease(32, 33). The cornerstones of NAFLD treatment are weight loss and increased amounts of exercise, but maintaining these lifestyle changes can be hard. Development of pharmacotherapy is in urgent needs when lifestyle modifications are not achieved and sustained(34). Currently, although many clinical trials are being investigated actively, there is still a lack of efficient drug for treatment of NASH. Our current work demonstrates the efficacy of givinostat for treatment of NASH via targeting inflammation and metabolism pathways.

As histone deacetylases could erase histone acetylation marks and regulate histone and chromatin structures, HDACs are classified into 4 groups: classI; classII; classIII known as the sirtuins, and classIV is represented by HDAC11 only (35, 36).Givinostat is a pan-HDAC inhibitor that belongs to hydroxamic acids family, with inhibition on classI, classII and classIII HDACs(37). Givinostat is being evaluated in phase III clinical trials at Italfarmaco company for the treatment of Duchenne’s muscular dystrophy(38), and is also undergoing Phase II clinical trials related to Myeloproliferative diseases or Polycythemia vera[Clinical trial NCT01901432]. Considering the crucial role of inflammation to NASH disease progression, we identified givinostat as a potent anti-inflammatory compound and assessed its pharmacologic efficacy for treatment of NASH. We found givinostat inhibited LPS and PA induced macrophage inflammatory activation *in vitro*, alleviated inflammatory cytokines expression and inflammatory cell infiltration in liver in NASH animal models. Our previous study also showed givinostat inhibited HSC activation and alleviated fibrogenesis in CCL4-induced liver fibrosis mouse model, which is consistent with other reports. The anti-inflammatory and anti-fibrogenesis role of givinostat has been revealed in several acute liver disease models, yet whether HDAC inhibitors could alleviated inflammation and fibrosis in chronic disease requires further study. Recent studies have demonstrated that givinostat could interfere inflammation and fibrosis progression in different diseases like intestinal epithelial inflammation or osteoarthritis(39, 40). This prompted us to examine whether givinostat could serve as a potential candidate for treatment of NASH, which remains undetermined up to now. Our study employed two models of NASH, MCD and FPC diets, to study the pharmacological potential of givinostat for treatment of NASH. MCD diet induced liver inflammation and fibrosis but lacks certain characteristics of human NASH pathology (like obesity or insulin resistance). FPC diet induced a phenotype of obesity, steatosis and steatohepatitis, resembling human NAFLD pathology, but lacking severe fibrosis. So we use both models and demonstrate the beneficial effects of givinostat for ameliorating hepatic steatosis, inflammation and fibrosis.

The RNA-seq and mechanistic data suggest givinostat regulated pathways related to steatosis, inflammation and fibrosis. Its multiple involvements in pathways protecting against NASH, may imply pleiotropic aspects of givinostat. The pathogenesis of NASH is complex and includes a complicated molecular network. Strategies targeting a single factor might not lead to a satisfying outcome(41). Drugs targeting different steps of NASH pathogenesis has been developed and tested in clinical trial. Lipid-lowering agents like Stearoyl-CoA Desaturase (SCD1) inhibitors(42) or Thyroid Hormone Receptor agonists(43) are aimed at modulating metabolism and lipid steatohepatitis. In addition to these, antidiabetic agents and antiobesity agents are also promising candidates(44, 45). Considering the progression of NAFLD and NASH, there are some other agents involved in anti-inflammatory and antifibrotic pathways. Among these, targeting Farnesoid X Receptor (FXR)(46), Obeticholic acid is in pre-registered condition. Drugs targeting chemokine receptors(47) and ASK1 are also being investigated(48). The current status of various clinical pipelines is consistent with complex pathogenesis of NAFLD and NASH. However, it is inconclusive that therapeutic treatment should intervene with which step. Drugs targeting single step or specific targets of NASH turn out to be disappointing in clinic, thus it is proposed that combinatory therapies targeting multiple disease drivers might result in more satisfying results. Givinostat achieves its therapeutic potential for NASH by simultaneously modulating lipid metabolic dysfunction while decreasing inflammation and fibrosis, making it an attractive potential drug for treatment of NASH but also a potential complementary partner for novel combination therapies.

Beyond Givinostat’s anti-inflammation and anti-fibrogenesis effect, transcriptomic analysis revealed that givinostat significantly reduced expression of diverse metabolic genes. KEGG pathway enrichment analysis revealed that the major hepatic transcriptional signature affected by givinostat was associated with lipid metabolism pathways like PPAR signaling pathway or lipid biosynthesis pathways. In support of our findings, studies in hepatic cell *in vitro* suggested that givinostat could diminish PA induced intracellular lipid accumulation. In past reports, inhibiting class I HDACs could prompt adipocytes differentiation to oxidative phenotype by regulating *Pparg* and *Ucp1* genes(49). Meanwhile, report shows that through disruption of the class IIa HDAC corepressor complex, whole-body energy expenditure and lipid oxidation could be increased(50). Our findings together with these clues indicate that epigenetic mechanism may participate in regulation of metabolism especially lipid homeostasis. Moreover, Bromodomain extraterminal domain (BET) proteins, another important epigenetic targets which are relative to histone modification reading, have been reported to be involved in lipid biosynthesis, uptake and intracellular trafficking(51). These indicated that drugs targeting epigenetic proteins could share extended indications in treating metabolic disorder diseases.

In summary, givinostat potently alleviated Diet-induced hepatic steatosis, inflammation as well as liver injury and fibrosis. Mechanically, givinostat inhibited expression of pathological pro-inflammatory genes as well as regulated a broad set of gene related to lipid metabolism. Givinostat could also directly inhibit HSCs activation and expression of ECM proteins *in vitro*. NASH is a complex, multifactorial disease, and is driven by multiple mechanisms including hepatic lipo-toxicity, inflammation, and fibrosis. Givinostat alleviated NASH by targeting multiple aspects crucial for NASH progression, suggesting the repurposing givinostat for treatment of NASH.

## Acknowledgments

We are extremely grateful to the National Centre for Protein Science Shanghai (Shanghai Science Research Center, Protein Expression and Purification system) for their instrument support and technical assistance. We gratefully acknowledge financial support from the National Natural Science Foundation of China (81070344 to G.L., 81803554 to Y.Z., 91853205, 81625022 and 81821005 to C.L.,), the Ministry of Science and Technology of China (2015CB910304 to Y.Z.) and National Science & Technology Major Project of China (2018ZX09711002 to Y.Z.)

## Author Contributions

H.H and S.F. performed immunofluorescence, Western-blot analysis, NASH experiments, corresponding data analysis and wrote the manuscript. X.Z. and S.F. contributed to manuscript writing and modifying, and analyzed the RNA-seq data. G.L., C.L. and Y.Z. conceived and supervised the project and revised the manuscripts.

